# Skeletal muscle fibre type determines mitochondrial and metabolic responses to hypoxia and pulmonary inflammation

**DOI:** 10.1101/2025.10.06.680623

**Authors:** Angelos Gavrielatos, Cécile Cottet-Rousselle, Noé Brocker, Cindy Tellier, Amel Achouri, Alexandre Prola, Hervé Dubouchaud, Clovis Chabert

## Abstract

**Background:** Chronic obstructive pulmonary disease (COPD) patients often experience skeletal muscle dysfunction that may result from a complex combination of mitochondrial dysfunction, metabolic reprogramming and fibre type transitions. Among other factors, pulmonary inflammation and hypoxia contribute to the COPD-associated muscle defects. Nevertheless, the precise molecular mechanisms and their effects across muscles with distinct metabolic profiles remain elusive. This study investigated the independent and combined effects of chronic pulmonary inflammation and chronic hypoxia on mitochondrial function, metabolic enzyme activity and fibre type composition in oxidative (soleus) and glycolytic (plantaris) muscles.

**Methods:** Adult male Wistar rats were subjected to chronic hypoxia (FiO_2_=10%) and/or chronic pulmonary inflammation (induced by bi-weekly intratracheal instillations of lipopolysaccharides). After 4 weeks of exposure, mitochondria from both soleus and plantaris were isolated to measure oxygen consumption rates, reactive oxygen species (ROS) emission, calcium retention capacity (CRC), respiratory complex activities and fatty acid oxidation capacity. Cross-sections from both muscles were analysed for fibre typology, fibre cross-sectional area (fCSA) as well as succinate dehydrogenase (SDH) and glycerol-3-phosphate dehydrogenase (GPDH) activities.

**Results:** Chronic hypoxia led to a decline in adenosine diphosphate-stimulated complex I (CI) respiration (p<0.01), Hydroxyacyl-Coenzyme A dehydrogenase activity (p<0.01), weight (p<0.05) and fCSA of type IIb fibres (p < 0.05) in plantaris. In contrast, chronic hypoxia increased CI-derived ROS emission (p<0.01) without changes in mitochondrial respiration or mass of soleus. No alterations in fibre typology were noticed in either muscle following the hypoxic exposure. Chronic pulmonary inflammation caused a reduction in mitochondrial CRC and an increase in GPDH activity in type IIa fibres of soleus (p<0.001) without any changes in fibre type distribution. Conversely, chronic pulmonary inflammation induced a downregulation of GPDH activity in plantaris type I and type IIa fibres, in parallel with an elevation in the SDH/GPDH ratio across all fibre types and a rise in the proportion of type IIx fibres.

**Conclusions:** Our results demonstrate fundamental differences in the responses to chronic hypoxia and chronic pulmonary inflammation between soleus and plantaris. While hypoxia affects predominantly the mitochondrial function and mass of plantaris, pulmonary inflammation drives metabolic reprogramming in both muscles that opposes their intrinsic functional specialisation. Additionally, soleus appears more vulnerable to permeability transition pore opening following pulmonary inflammation. Notably, these mitochondrial alterations seem to occur independently of fibre type shifts highlighting the central role of intrinsic mitochondrial maladaptations in the COPD-associated muscle dysfunction.

## 1. Introduction

Skeletal muscle dysfunction and atrophy are common features in a range of pathological conditions including chronic obstructive pulmonary disease (COPD). Notably, more than 30% of COPD patients experience skeletal muscle impairments which serve as critical indicators of deteriorated quality of life, exercise intolerance and increased reliance on healthcare resources (1). To date, accumulating evidence confirms that limb muscles in COPD may exhibit alterations in mitochondrial function and structure including a reduction in mitochondrial density (2), a depressed respiratory capacity (3) and a decline in oxidative enzyme activity (4,5). This metabolic reprogramming may further be associated with a faster opening of the mitochondrial permeability transition pore (mPTP) with concomitant cytochrome c release (6) and an elevated reactive oxygen species (ROS) emission (3), collectively pointing to a profound mitochondrial remodelling within the COPD-affected muscle.

The pathogenesis of structural and functional alterations in skeletal muscle of COPD patients is multifactorial involving oxidative stress, pulmonary inflammation, alveolar hypoxia, malnutrition, physical inactivity, and corticosteroid administration (7,8) However, their individual contributions to skeletal muscle mitochondrial alterations remain unclear. The pulmonary origin of COPD establishes the lungs as the primary driver of the skeletal muscle mitochondrial pathology likely through chronic airway inflammation, which may propagate to the muscle via spill-over of inflammatory mediators (9), and alveolar hypoxia which may lead to muscle tissue hypoxia and capillary rarefaction (10,11) initiating peripheral muscle wasting before secondary confounders (e.g., physical inactivity) exacerbate it. Chronic hypoxia has been more extensively investigated for its impact on skeletal muscle mitochondria, with the available data suggesting an elevation in ROS production (12), a decrease in mitochondrial respiration (13) and a shift towards glycolytic metabolism (14). Yet, distinguishing these effects from the compounding influence of physical inactivity remains challenging, as many studies may have examined hypoxia concurrently with inactivity. Furthermore, variation in parameters of hypoxic exposure (e.g., continuous vs intermittent, duration, fraction of inspired O_2_ (FiO_2_)) complicates cross-study comparisons (15). In contrast, despite the well-studied role of inflammation in muscle homeostasis (16), the consequences of chronic pulmonary inflammation on skeletal muscle mitochondria are poorly defined with current evidences largely confined to acute models.

We have previously demonstrated that chronic pulmonary inflammation, blunts the overload-induced hypertrophy in oxidative rat soleus (17) while De Theije and colleagues (18) have reported that 21 days of hypoxic exposure leads to atrophy, independently of the hypoxic hypophagia, in glycolytic mice EDL but not in soleus. Considering the divergent mitochondrial signatures between oxidative and glycolytic muscles, especially with respect to respiratory properties, ROS metabolism and mPTP regulation by Ca^2+^ (19), mitochondrial responses to inflammation and hypoxia may also present a fibre type-specific heterogeneity. Nevertheless, neither study assessed mitochondrial function or metabolic parameters under these conditions. Therefore, elucidating the differential susceptibility of glycolytic and oxidative muscles to these stressors remains imperative for uncovering mechanisms underlying compartmentalised muscle dysfunction and atrophy while guiding the development of phenotype-specific rehabilitation, personalised anti-atrophy interventions and biomarker discovery.

Facing these limitations, the aim of the current study was to unravel the isolated and combined effects of chronic pulmonary inflammation and chronic hypoxia on oxidative soleus and glycolytic-leaning plantaris muscles of adult male rats with a focus on mitochondrial function and key metabolic enzymes. Given that fibre type transitions are commonly observed in skeletal muscle of COPD patients (20), we also performed a fibre typology analysis in both muscles to explore whether any mitochondrial changes are accompanied by alterations in the myosin heavy chain (MHC) phenotype. Based on prior findings from our laboratory, we hypothesised that the highly oxidative soleus, with its dense capillary network, would exhibit greater sensitivity to pulmonary inflammation due to enhanced perfusion of systemic inflammatory mediators, whereas the glycolytic-leaning plantaris, with sparser vascularisation, would develop more pronounced pathological alterations under chronic hypoxia.

## 2. Methods

### 2.1. Animals

The study was conducted in accordance with the guidelines of the European Union Directive (2010/63/EU) on laboratory animal care and use and received approval from the local Ethics Committee (Cometh 12) affiliated with the laboratory and authorised by the French Ministry of Higher Education, Research and Innovation (APAFIS #40584-2023013114463378). Twenty-eight adult male Wistar rats aged 20 to 27 weeks with mean body weight ± SD of 530 ± 41 g were housed in individual cages with free access to food (ref: 3430; SAFE, Augy, France) and water. Body mass and food intake were weekly recorded.

Rats were assigned to one of the following groups for 28 days until sacrifice: 1) Normoxia control (NC), 2) Chronic hypoxia (H), 3) Chronic pulmonary inflammation (I), and 4) H + I (HI). An overview of the experimental design is provided in Figure 1. Chronic pulmonary inflammation was induced by intratracheal instillations of lipopolysaccharides (LPS) at 1 mg/kg of body weight (E.coli, serotype O55:B5; Sigma-Aldrich Chemical Co. ™, St Louis, Missouri, USA) in NaCl 0.9% twice per week for 4 weeks after isoflurane anaesthesia. The final LPS instillation occurred ≥ 60 hours prior to sacrifice to preclude confounding effects by acute-phase inflammatory responses. Animals from the H and HI groups were exposed for 28 days to normobaric hypoxia in O_2_-controlled chambers (FiO_2_: 10%). The hypoxic chambers were opened for 10-15 min each day to weigh the animals, the food pellets and to perform LPS instillations to the rats from the HI group. At the end of the study, animals were euthanised by exsanguination from the abdominal aorta following intraperitoneal injection of ketamine and xylazine (80 mg/kg and 50 mg/kg of body weight, respectively). Soleus and plantaris were bilaterally dissected, with right hindlimb muscles processed for isolation of mitochondria while left hindlimb muscles were cryopreserved in liquid nitrogen-cooled isopentane, and stored at -80°C until subsequent cryosectioning.

**Fig.1.**
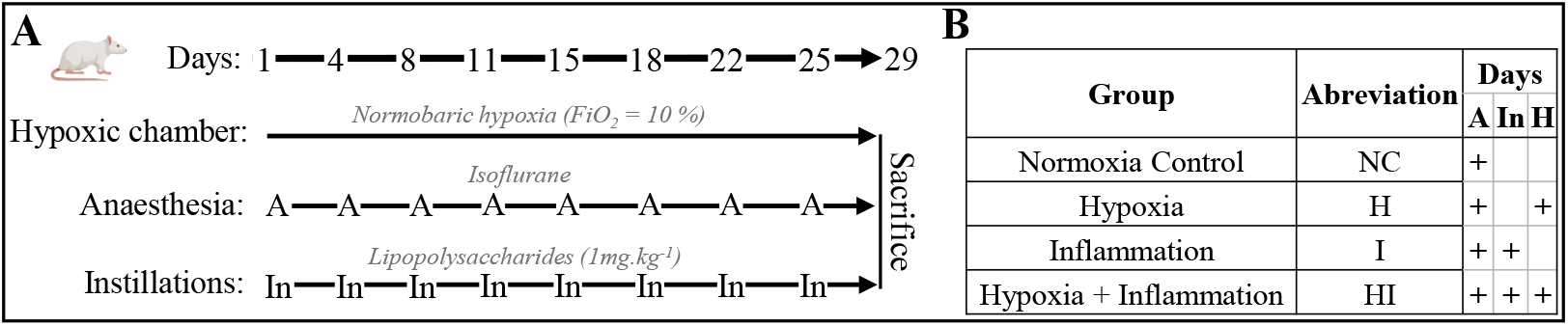
Experimental design of the study. **A**: Timeline of the different procedures carried out. **B**: summary table of the procedures performed on animals according to their group. A: Anaesthesia; In: Intratracheal instillations.

### 2.2. Mitochondria isolation

Immediately after dissection, soleus and plantaris were immersed in ice-cold isolation buffer (150 mM saccharose, 75 mM KCl, 50 mM Tris Base, 1 mM KH_2_PO_4_, 5 mM MgCl_2_, 1 mM EGTA, pH 7.4) and were trimmed of connective tissue before transfer to isolation buffer supplemented with 0.2% fat-free bovine serum albumin (BSA) at 4°C. Muscles were minced with scissors and incubated with subtilisin (0.24 mg/ml) for 1 min before addition of isolation buffer and 0.2% BSA to stop the reaction. The suspension was subsequently homogenised using a motor-operated glass-Teflon pestle homogeniser and centrifuged twice at 800 g for 10 min at 4°C to remove cell debris and nuclei. Mitochondria from the resulting supernatant were pelleted by centrifuging at 8000 g for 10 min at 4°C. The mitochondrial pellet was resuspended in 0.2 ml respiration buffer (125 mM KCl, 20 mM Tris Base, 1mM EGTA, pH 7.2) and kept on ice until use. The supernatant containing the cytosolic fraction was also collected and stored at -80C until spectrophotometric analysis of lactate dehydrogenase activity (LDH). Mitochondrial protein concentration was determined using the bicinchoninic acid assay (BCA; Pierce™, ThermoFisher Scientific, Waltham, MA, USA) with BSA standards (Pierce™).

### 2.3. Oxygen consumption measurement

Mitochondrial oxygen consumption was measured using a Clark-type O_2_ electrode (Oxygraph+, Hansatech Instruments, Pentney, UK). Mitochondria (0.2 mg/ml) were incubated in respiration buffer supplemented with 0.15 % BSA and 0.1 M inorganic phosphate (Pi) at 30°C under constant electromagnetic stirring. Mitochondria were energised using CI (glutamate/malate, GM; 5 mM/2.5 mM) and Complex II (CII) substrates (succinate, S; 5 mM) either alone or in combination (GMS). Oxygen consumption was recorded before (state 2) and after the addition of adenosine diphosphate (ADP) 1 mM (state 3) and following the addition of oligomycin (1 μg/ml) (state 4). When succinate was utilised as a substrate alone, rotenone (4 μM) was added to inhibit CI. The activity of cytochrome c oxidase (Complex IV, CIV) was determined by sequential additions of antimycin A (AA; 2 μM), Ascorbate (2.5 mM), TMPD/Ascorbate (0.5 mM/0.2 mM) and finally the uncoupler DiNitroPhenol (150 μM). Mitochondrial respiration rates were expressed as nmol O/min/mg mitochondrial protein.

### 2.4. Mitochondrial H_2_O_2_ release

Mitochondrial hydrogen peroxide (H_2_O_2_) release was quantified with a fluorimeter (F2500 Hitachi, Tokyo, Japan) by monitoring the linear increase in fluorescence due to the enzymatic oxidation of Amplex Red to resorufin (excitation, exc, at 560 nm; emission, em, at 584 nm) by H_2_O_2_ in the presence of horseradish peroxidase (HRP). Isolated mitochondria (0.2 mg/ml) were incubated at 30°C in mitochondrial respiration buffer, supplemented with 0.15 % BSA, 6 U/ml of HRP and 1 μM of Amplex Red under constant electromagnetic stirring. Mitochondrial H_2_O_2_ emission was assessed in basal conditions with GM (5 mM/2.5 mM) or S (5 mM) followed by sequential addition of rotenone (3 μM) to maximise CI-derived H_2_O_2_ (GM) or block reverse electron flow (S) and AA (3 μM), only with GM, to trigger complex III-derived H_2_O_2_ from both complexes (I + III). Results were expressed as pmol H_2_O_2_/min/mg mitochondrial protein using a known amount of H_2_O_2_ as a standard.

### 2.5. Calcium retention capacity

To assess the mPTP sensitivity, the mitochondrial calcium retention capacity (CRC) was quantified using a spectrofluorimeter (Horiba PTI QM-400 Special QuantaMaster, Irvine, California, USA). Mitochondria (0.1 mg/ml) were incubated at 30°C under continuous electromagnetic stirring in 250 mM saccharose, 10 mM Pi, 10 mM Tris-MOPS, 10 mM EGTA, pH 7.4, containing 125 μM calcium green-5N (Calcium Green™-5N, ThermoFisher Scientific) as the Ca^2+^-sensitive fluorescent probe. Mitochondria were energised with GM with or without the presence of 1 μM cyclosporin A (CsA) inhibitor of cyclophilin D (CyP-D). Ca^2+^pulses of either 1.5 nmol for soleus or 2 nmol for plantaris were added at 1-minute intervals. Mitochondrial CRC was calculated as the total amount of Ca^2+^ (nmol/mg protein) required to trigger opening of the mPTP, indicated by a sharp increase in fluorescence due to Ca^2+^ release from the mitochondrial matrix.

### 2.6. Respiratory chain complex I, HAD, and LDH activities

To determine CI activity, mitochondria (10 μg/ml) were incubated at 30°C in 47.5 mM KH_2_PO_4_, pH 7.5, containing 95 μM EDTA, 3.75 mg/ml BSA, 100 µM NADH, and 100 μM decylubiquinone. NADH oxidation kinetics was monitored at 340 nm for 4 min with or without the presence of 10 µM rotenone to calculate total and non-specific activities, respectively. To assess HAD activity, mitochondria (10 μg/ml) were incubated at 30°C in 47.5 mM KH_2_PO_4_, pH 7.5 supplemented with 200 µM NADH, 60 µM EDTA, and 40 mM imidazole. Following initial measurement of non-specific activity, total HAD activity was quantified by monitoring the decrease in the NADH absorbance at 340 nm after addition of acetoacetyl-CoA (3-ketoacyl-CoA analogue). LDH activity was determined in plantaris using the cytosolic fractions obtained during mitochondrial isolation. After determination of protein concentration using the BCA assay, samples were incubated at 37°C in 100 mM TRIS buffer, pH 7.1, 15 mM NADH. Following initial measurement of non-specific activity, total LDH activity was quantified by monitoring the decrease in the NADH absorbance at 340 nm after addition of pyruvate. CI, HAD and LDH specific activities were all expressed in µmol/min/mg protein.

### 2.7. Histochemistry and Immunofluorescence

Muscle cross-sections (8 µm) were cut at -20°C (Leica CM3050 S, Wetzlar, Germany) and mounted on SuperFrost Plus slides (Epredia, NH, USA). Slides were air-dried at room temperature (RT) and stored at -20°C. All the slides were stained within one month after cryosectioning.

Activity of succinate dehydrogenase (SDH) and glycerol-3-phosphate dehydrogenase (GPDH) were determined histochemically in different slides. Freshly prepared and pre-warmed at 37°C, either SDH (50 mM sodium succinate, 0.45 mM 1-methoxyphenanzine methosulfate, 5 mM sodium azide and 4.95 mM nitroblue tetrazolium chloride) or GPDH (50 mM Racemic-G1P, 1 mM menadione, 5 mM sodium azide and 4.95 mM nitroblue tetrazolium chloride) reaction medium, diluted in 0.1 M Tris-Maleate buffer (pH 7.5) with 10% polyvinyl alcohol, was applied to tissue sections. Enzyme reactions proceeded at 37°C for 10 min under light-protected conditions. Reaction was stopped by rinsing the sections with ice-cold PBS. The specificity of the staining was validated using competitive inhibitors: 50 mM dihydroxyacetone phosphate for GPDH and 250 mM malonate for SDH.

To determine muscle fibre typology and fibre cross-sectional area (fCSA), slides were subsequently incubated for 1 h at 37°C with primary antibodies targeting MHC isoforms diluted in PBS, 1% goat serum and 2% BSA. Soleus samples were immunolabelled with antibodies against MHC I (BA-F8 IgG2b, 1:50) and MHC IIa (SC-71 IgG1, 1:100) isoforms. Plantaris specimens were incubated with antibodies against MHC I (BA-F8, 1:100), MHC IIa (SC-71, 1:200), and MHC IIb (BF-F3 IgM, 1:100) while the MHC IIx isoform was identified by exclusion. All the antibodies were obtained from the Developmental Studies Hybridoma Bank (University of Iowa, IA, USA). Species- and subclass-specific fluorescent secondary antibodies, purchased from Invitrogen (Waltham, MA, USA) were then incubated with slides for 1h at 37°C (Alexa Fluor 488, ref: A21141, 1:300 for BA-F8; Alexa Fluor 568, ref: A21124, 1:300, for SC-71; Alexa Fluor 647, ref: A21238, 1:300, for BF-F3). Following subsequent PBS washes, cellular membranes were stained with CF®405S WGA (1:200 CliniSciences, Nanterre, France) for 10 minutes at RT. Slides were air-dried, mounted with ProLong Gold Antifade (Invitrogen), coverslipped, and cured overnight at RT under light-protected conditions.

### 2.8. Image acquisition and analysis

Representative images illustrating MHC, WGA, GPDH and SDH staining are provided in supplementary Figure S1. Images were acquired within 24 h post-mounting using a Leica DMi8 wide-field epifluorescence microscope (Leica Microsystems, Wetzlar, Germany) equipped with a 20x/0.8 dry objective and a camera (Leica DFC 9000GT). Raw data collection and processing were done using the manufacturer’s LAS-X software. Fluorescence signal was recorded with four filter cubes (CF®405S WGA: DAPI cube, λexc = 390 nm, λem = 435 nm; Alexa Fluor 488: 3-GFP cube, λexc = 475 nm, λem = 519 nm; Alexa Fluor 568: 5-Rhodamine cube, λexc = 555 nm, λem = 594 nm; Alexa Fluor 647, 7-Cy5 cube, λexc = 635 nm, λem = 641 nm). SDH and GPDH enzyme activities were imaged using transmitted light microscopy (brightfield mode) on the same Leica DMi8 microscope (illumination intensity: 100%). All morphological analyses and subsequent quantification were conducted using Fiji (v1.54m), an enhanced distribution of ImageJ (National Institute of Health, USA) by two investigators blinded to experimental group coding.

To determine fibre type classification, an assisted automatic detection of the fibres based on WGA membranes staining was performed and the mean signal intensity in each channel was measured. For each channel, a threshold was established, beyond which fibers were classified as positive for measured MHC isoform. Fibre typology was determined by manually counting all fibres within entire muscle cross-sections. The abundance of each fibre type was calculated as a percentage of the total fibre count. Fibre cross-sectional areas were determined by encircling WGA-stained cell membranes and analysing a minimum of 50 fibres per type except for the soleus type IIa fibres. To determine the mean fCSA of each group, mean fCSA of each sample were averaged. For the SDH and GPDH analyses, the enzymatic activity was calculated by measuring the integral density (sum of pixel values in a defined area) of each fibre divided over the fCSA and expressed in arbitrary units (AU)/min. Group-level mean activities for SDH and GPDH were derived by averaging sample means.

### 2.9. Statistical analysis

Normality of the data was examined using the Shapiro-Wilk test and visual inspection of histograms and boxplots. Non-normal distributions were normalised with log_10_ or square root transformations. To examine the effect of the two experimental conditions (chronic constant hypoxia and chronic pulmonary inflammation), a 2-way analysis of variance (ANOVA) was performed with each of the experimental conditions (condition 1: normoxia or hypoxia; condition 2: with or without pulmonary inflammation) as between-subjects factors. Post-hoc analyses for significant interaction effects employed Bonferroni-corrected one-way ANOVAs and independent t-tests. All fibre typology variables were treated as a family of hypotheses due to their inherent interdependence and were analysed using a 2-way multivariate analysis of variance (MANOVA). Model assumptions were evaluated via Box’s M test and visual examination of correlation matrices. Results are presented as mean ± standard error of mean (SEM). A threshold of p < 0.05 was used to define statistical significance. Data points deviating ≥ 2 SD from the group mean were excluded from the data as outliers. Effect sizes were quantified by partial eta square (η^2^_p_) values, with a η^2^_p_ value ≥ 0.14 indicating a large effect. All the p-values ranging between 0.05 and 0.06 were interpreted as statistical trends indicating marginal significance. Statistical analyses were conducted with SPSS (v27, IBM, Chicago, IL, USA) and all figures were generated with GraphPad Prism (v8.0.2 GraphPad Software, San Diego, USA) except for Figure 1 that was designed using BioRender.com.

## 3. Results

### 3.1. Body weight, food intake and muscle weight

Starting body weight was the same among the groups (Figure 2A). However, at the day of sacrifice, final body weight was lower in the chronic hypoxia condition compared to normoxic conditions (NC + I: 549 ± 39 g. vs H + HI: 454 ± 44.4 g; p < 0.001; η^2^_p_ = 0.64; Figure 2A). That difference is associated with a significant decrease of body weight during the first week of exposure (−14.5%; p < 0.001; η^2^_p_ = 0.82; Figure 2B). Body weights were unchanged over the last 3 weeks of exposure in all the groups (Figure 2B). Cumulated food consumption over the study period was lower in animals exposed to hypoxia (NC + I: 716 ± 67.2 g vs H + HI: 447 ± 61.5 g; p < 0.001; η^2^_p_ = 0.85; Figure 2C). When adjusted to body mass, weekly food consumption was lower in hypoxia groups compared to normoxia groups during the first three weeks of exposure but not during the final week (week 1: - 92%, p < 0.001, η^2^_p_ = 0.82; week 2: -58%, p < 0.001, η^2^_p_ = 0.82; week 3: -33%, p < 0.001, η^2^_p_ = 0.39; week 4: -14%, p = 0.132, η^2^_p_ = 0.096; Figure 2D). Hypoxia exposure induced a decline in plantaris muscle weight (NC + I: 0.52 ± 0.06 g vs H + HI: 0.45 ± 0.07 g; p < 0.05; η^2^_p_ = 0.26; Figure 2F) but not in soleus muscle, when compared to normoxia conditions.

**Fig.2.**
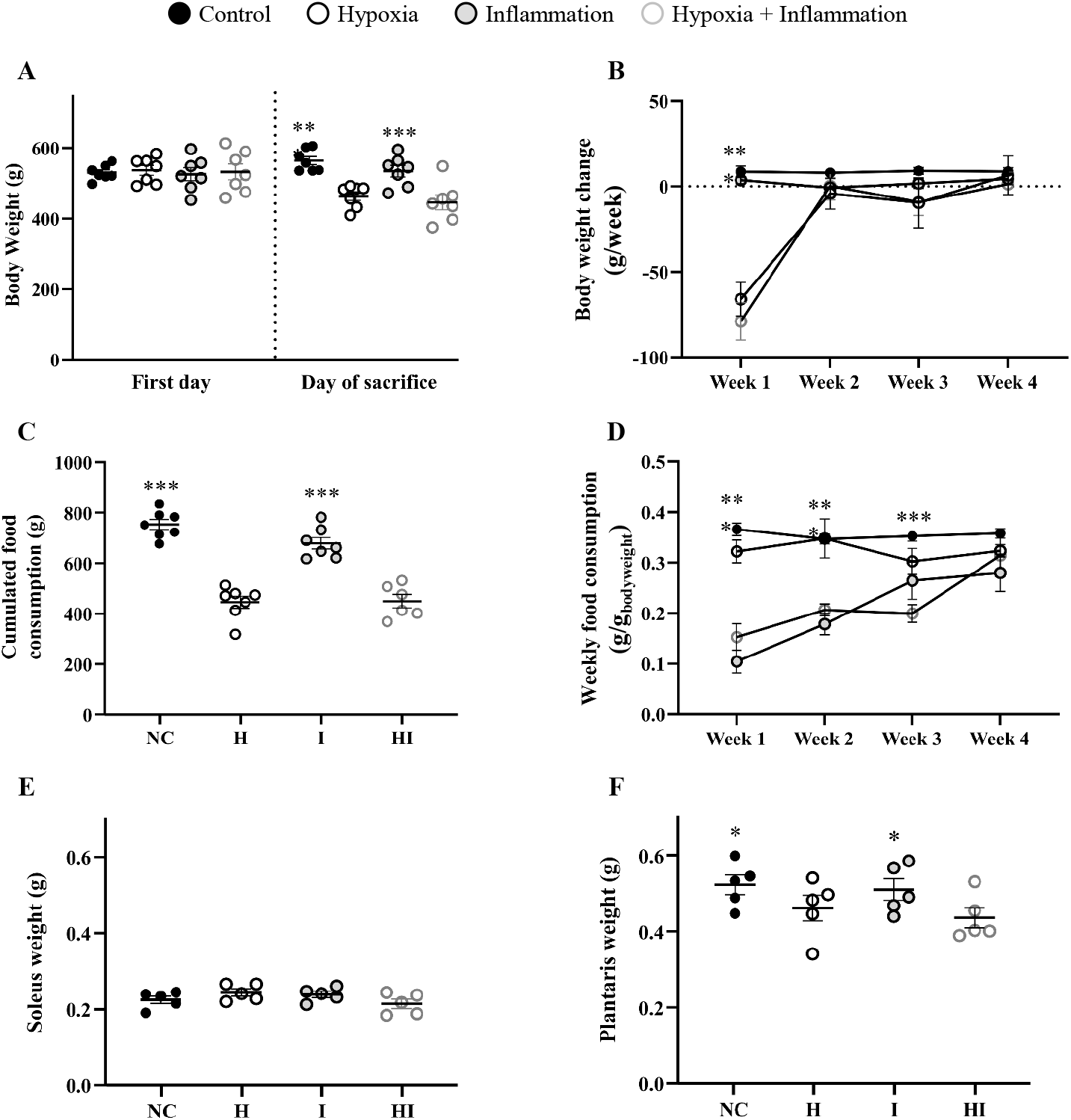
Body mass, food intake and muscle weight under hypoxia and pulmonary inflammation conditions. **A**: Initial and final body weight. **B**: Weekly changes in body weight throughout the 4-week exposure. **C**: Total cumulated food intake over the 4-week study period. **D**: Weekly changes in the relative food intake of the rats throughout the 4-week study period. **E**: Soleus weight at sacrifice. **F**: Plantaris weight at sacrifice. Circles indicate individual data points and lines represent means ± SEM. ***: p < 0.001, global effect of hypoxia (NC + I vs H + HI). *: p < 0.05, global effect of hypoxia (NC + I vs H + HI).

### 3.2. Mitochondrial respiration

Neither condition changed the respiration under basal (substrate alone), state 3 (ADP-stimulated) or state 4 (non-phosphorylating) conditions in soleus mitochondria, regardless of the substrates fuelling the electron transport chain (Figure 3A-C). Similarly, the respiratory control ratio (RCR: state 3/state 4) was not affected by the exposure to pulmonary inflammation and/or hypoxia in soleus under any condition (supplementary Figure S2A). Soleus CIV respiration appeared also unaffected (supplementary Figure S3A) by inflammation and/or hypoxia. In contrast, chronic hypoxia led to a decline in state 3 respiration with complex I substrates in mitochondria from plantaris muscle (−45%, p < 0.01; η^2^_p_ = 0.29; Figure 3D). However, neither hypoxia nor inflammation altered state 2 or state 4 CI respiration (Figure 3D) or RCR (supplementary Figure S2B) in mitochondria from plantaris. Furthermore, no changes in plantaris mitochondrial respiration or RCR was observed for any other conditions (Figure 3E-F, supplementary Figures S2B and S3B).

**Fig.3.**
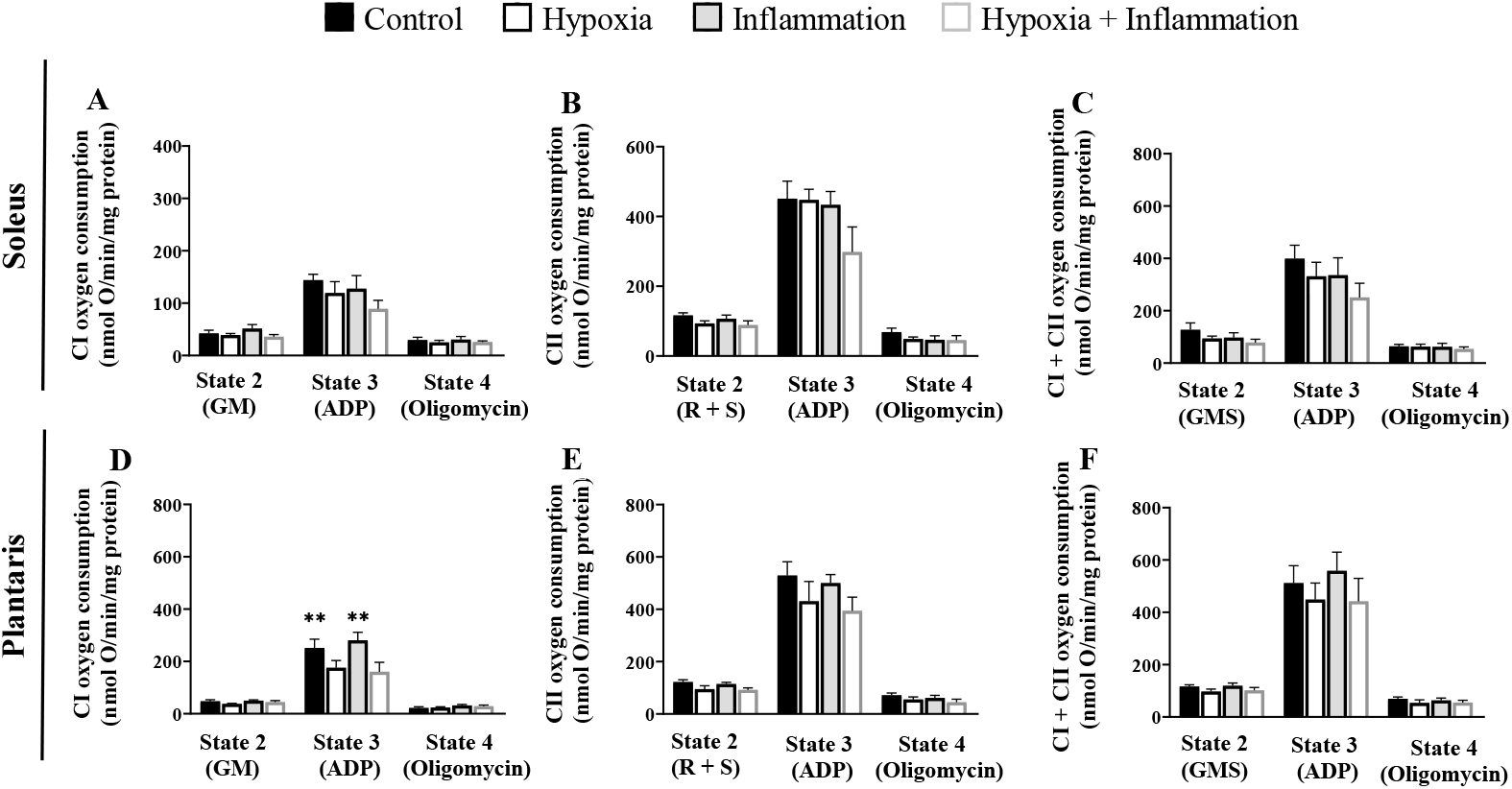
Oxygen consumption in isolated mitochondria from soleus and plantaris muscle after exposure to hypoxia and inflammation. Soleus and plantaris isolated mitochondrial oxygen consumption measured by oxygraphy with Glutamate + Malate (**A, D**), Succinate + Rotenone (**B, E**) and Glutamate + Malate + Succinate (**C, F**) substrates (basal state 2 condition), after ADP addition (ADP-stimulated State 3 condition) or after addition of oligomycin (non-phosphorylating State 4 condition). Data are shown as the mean ± SEM. Soleus data are presented in panels **A-C** while plantaris data are shown in panels **D-F**. **: p < 0.01, global effect of hypoxia (NC + I vs H + HI).

### 3.3. Mitochondrial H_2_O_2_ release

Basal mitochondrial H_2_O_2_ release, as indicator of mitochondrial ROS production, was significantly higher in soleus isolated mitochondria after exposure to chronic hypoxia with GM as substrate (+ 54.8%, p < 0.01, η^2^_p_ = 0.39; Figure 4A). Consistently, hypoxia exposure was associated with a higher ratio of CI-linked H_2_O_2_ emission over basal CI respiration (+ 57.6%, p < 0.001; η^2^_p_ = 0.43; supplementary Figure S4A). However, this effect in soleus muscle was abolished when rotenone was used to block CI or when AA was added (Figure 4A). Notably, neither hypoxia nor pulmonary inflammation altered H_2_O_2_ emission in soleus isolated mitochondria, whether measured during combined forward and reverse electron transfer pathways using S or isolated reverse electron flow revealed by CI inhibition with rotenone (Figure 4B). These findings were consistent with the lack of changes in ROS release with succinate as substrate when normalised to CII basal respiration rate (supplementary Figure S4A). In plantaris, hypoxia and inflammation exposures did not have any effect on isolated mitochondria H_2_O_2_ release regardless the combination of substrates and inhibitors (Figure 4D-F, supplementary Figure S4B).

**Fig.4.**
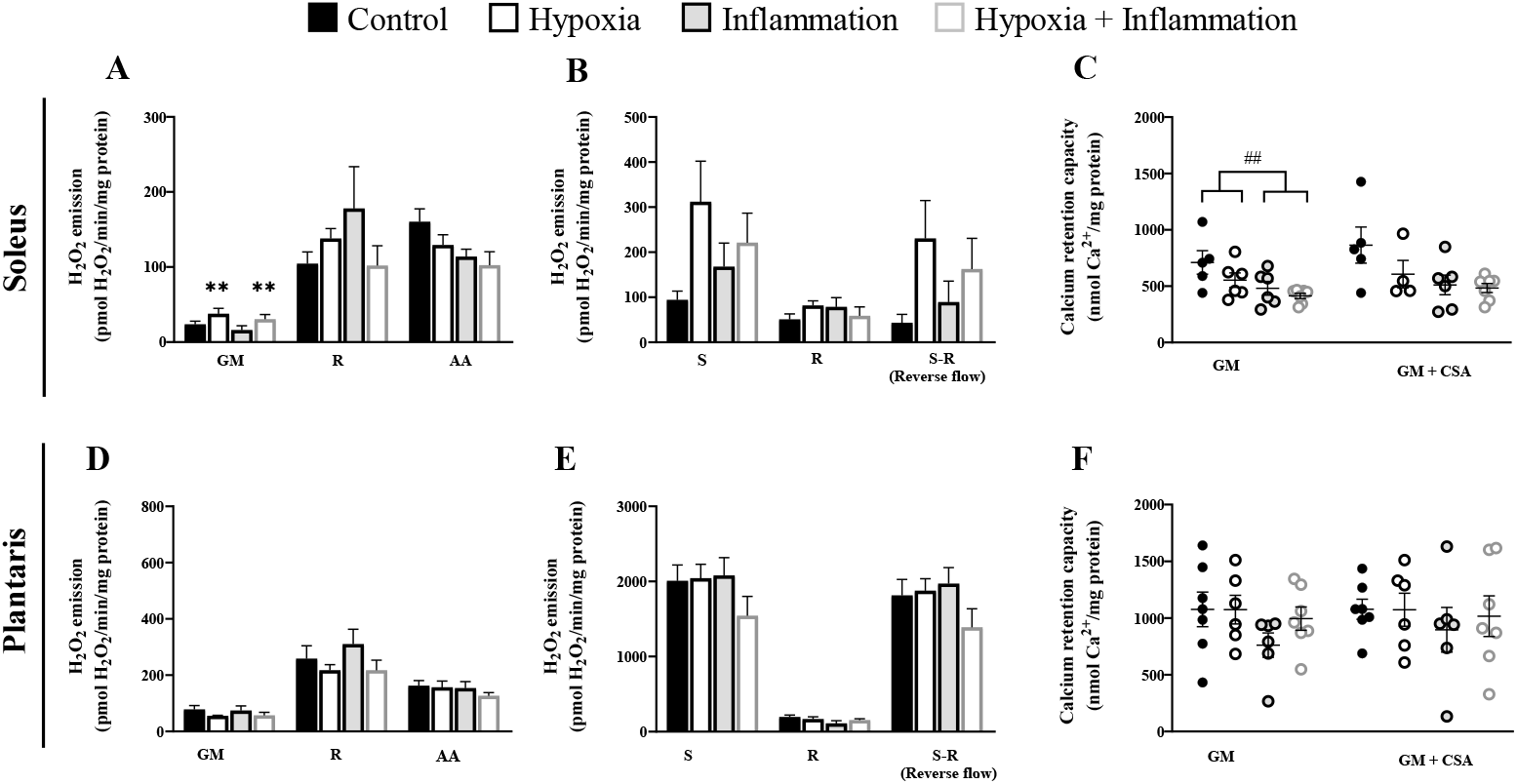
Effect of hypoxia and inflammation exposures on isolated mitochondrial H_2_O_2_ emission and calcium retention capacity. Soleus (**A, B**) and plantaris (**D, E**) isolated mitochondrial H_2_O_2_ release with substrate alone and after rotenone and AA inhibition. Data are shown as the mean ± SEM. C, f: Soleus (**C**) and plantaris (**F**) isolated mitochondrial CRC was measured with Glutamate + Malate or with Glutamate + Malate + Cyclosporin A. Circles indicate individual data points and lines represent means ± SEM. **##:** p < 0.01, global effect of pulmonary inflammation (NC + H vs I + HI).

### 3.4. Calcium retention capacity

Chronic pulmonary inflammation led to a decline in soleus mitochondrial CRC when GM was used to fuel the electron transport chain (NC + H: 624 ± 202 nmol Ca^2+^/mg prot vs I + HI: 445 ± 112 nmol Ca^2+^/mg prot; p < 0.001; η^2^_p_ = 0.3; Figure 4C). However, inhibition of the mitochondrial permeability transition pore with CsA reversed this effect of inflammation exposure (Figure 4C). No change in plantaris CRC was detected whether mitochondria were incubated with GM alone or GM combined with CsA (Figure 4F).

### 3.5. CI, HAD, and LDH enzymatic activities in isolated mitochondria

There were no effect of pulmonary inflammation or hypoxia on HAD activity of soleus mitochondria (Figure 5A). However, exposure to chronic hypoxia led a significant decrease in HAD activity of plantaris mitochondria (−31.5%; p < 0.01; η^2^_p_ = 0.32; Figure 5C). Neither environmental constraint induced alterations in isolated CI activity in soleus or plantaris muscles (Figure 5B & D).

**Fig.5.**
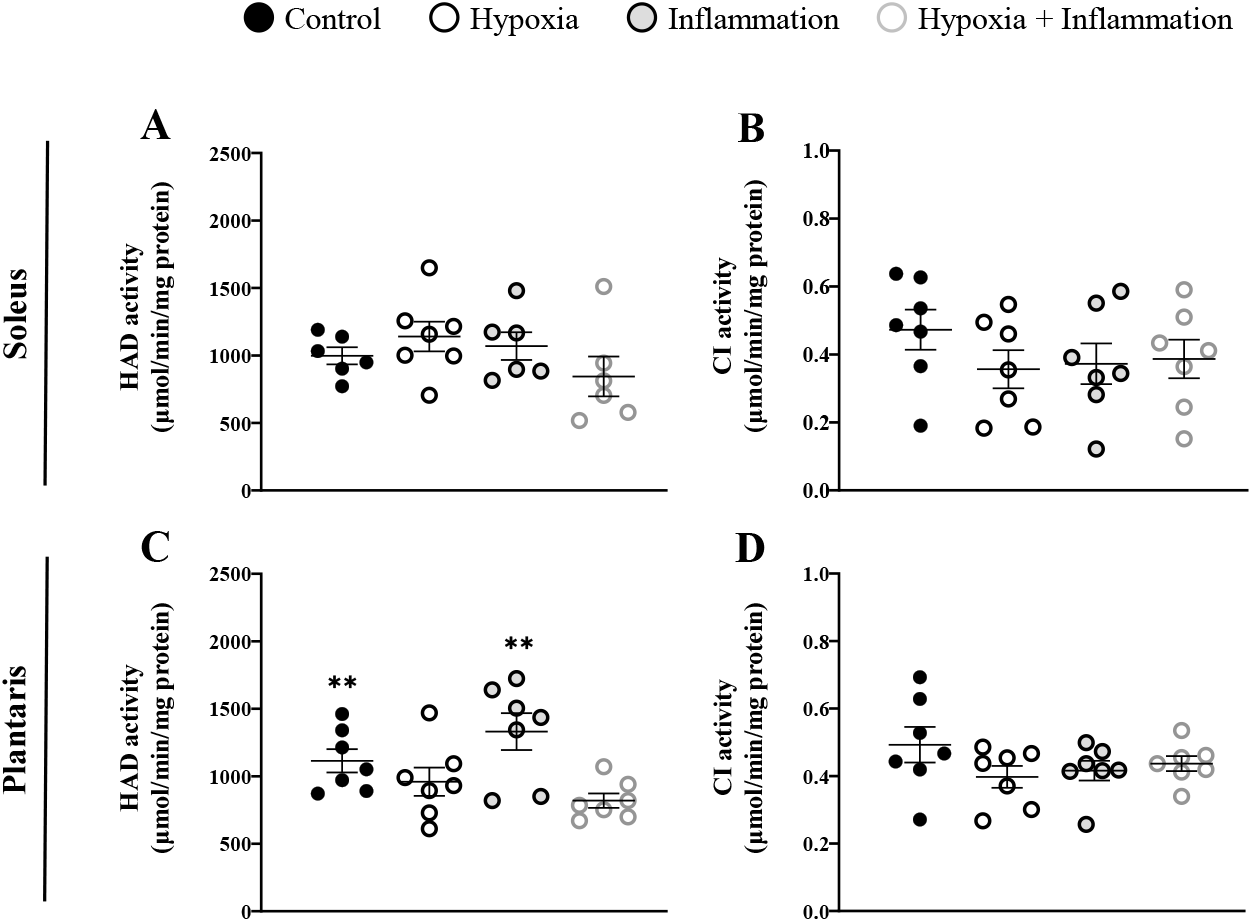
Effect of hypoxia and inflammation exposure on HAD activity and CI activity. HAD activity was measured in isolated mitochondria of soleus (**A**) and plantaris (**C**). CI activity was measured in isolated mitochondria of soleus (**B**) and plantaris (**D**). Circles indicate individual data points and lines represent means ± SEM. **: p < 0.01, global effect of hypoxia (NC + I vs H + HI).

### 3.6. Fibre typology and fCSA

There were no alterations in the fibre type composition of soleus in response to either intervention (Figure 6A-B). In contrast, chronic pulmonary inflammation led to a significant elevation in the proportion of type IIx fibres in plantaris (NC + H: 41.6 ± 4.8 % vs I + HI: 46.6 ± 4 %, p < 0.05, η^2^_p_ = 0.26, Figure 6C). Furthermore, exposure to either experimental condition did not elicit any changes in the hybrid fibres of plantaris (Figure 6D). Neither conditions affected the type I or type IIa fCSA in soleus muscle (supplementary Figure S5A-B). However, in plantaris, hypoxia led to a decrease in the fCSA of type IIb fibres but not in type I, IIa or IIx (type IIb: NC + I: 5332 ± 1182 μm^2^ vs H + HI: 4464 ± 591; p < 0.05, η^2^_p_ = 0.22, supplementary Figure S5C-F).

**Fig.6.**
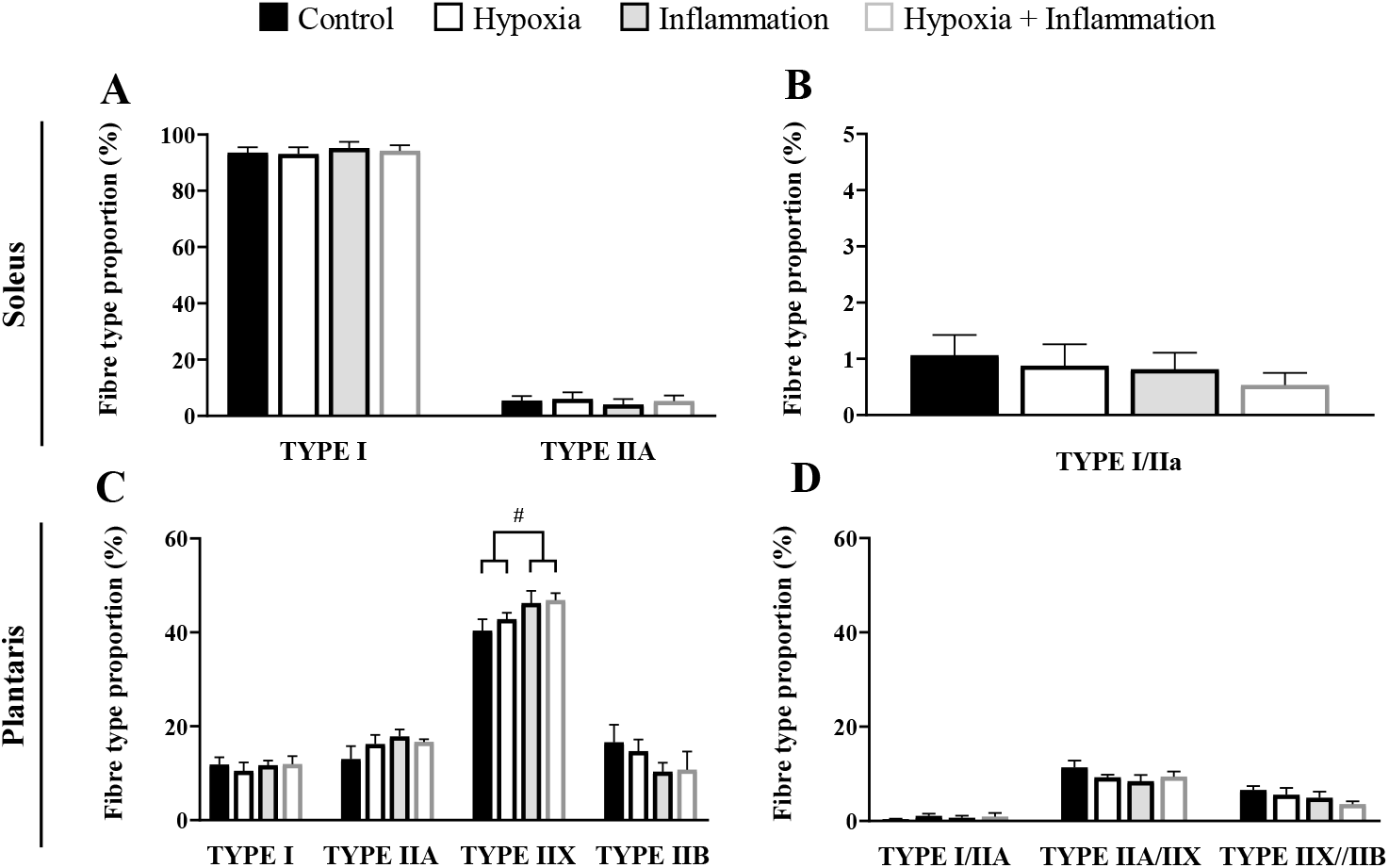
Effects of hypoxia and inflammation exposure on fibre typology. Percentage of pure and hybrid fibres in soleus (**A-B**) and plantaris (**C-D**). Data are shown as means ± SEM. **#:** p < 0.05, global effect of pulmonary inflammation (NC + H vs I + HI).

### 3.7. SDH and GPDH activities

We did not observe any effect of pulmonary inflammation or hypoxia in the SDH activity of type I or type IIa fibres in soleus (Figure 7A-B). In addition, we report no differences in the GPDH activity (Figure 8A) or SDH/GPDH ratio (supplementary Figure S6A-B) in soleus type I fibres. In contrast, pulmonary inflammation increased the GPDH activity in soleus type IIa fibres (+ 15.9 %, p < 0.001, η^2^_p_ = 0.48, Figure 8B). Likewise, there was a tendency for a reduction in the SDH/GPDH ratio in type IIa fibres of soleus in the animals exposed to pulmonary inflammation (−32.9%, p = 0.051, η^2^_p_ = 0.26, supplementary Figure S6B). Neither environmental condition altered the plantaris SDH activity regardless the fibre type (Figure 7C-F). Conversely, pulmonary inflammation induced a significant decline in the GPDH activity in the type I and type IIa fibres of plantaris (type I: - 31.8%, p < 0.05, η^2^_p_ = 0.33, Figure 8C; type IIa: - 39.5%, p < 0.01, η^2^_p_ = 0.52, Figure 8D) but not in type IIx or type IIb (Figure 8E-F). Finally, pulmonary inflammation led to a significant increase in the SDH/GPDH ratio across all the fibre types of plantaris (supplementary Figure S6C-F).

**Fig.7.**
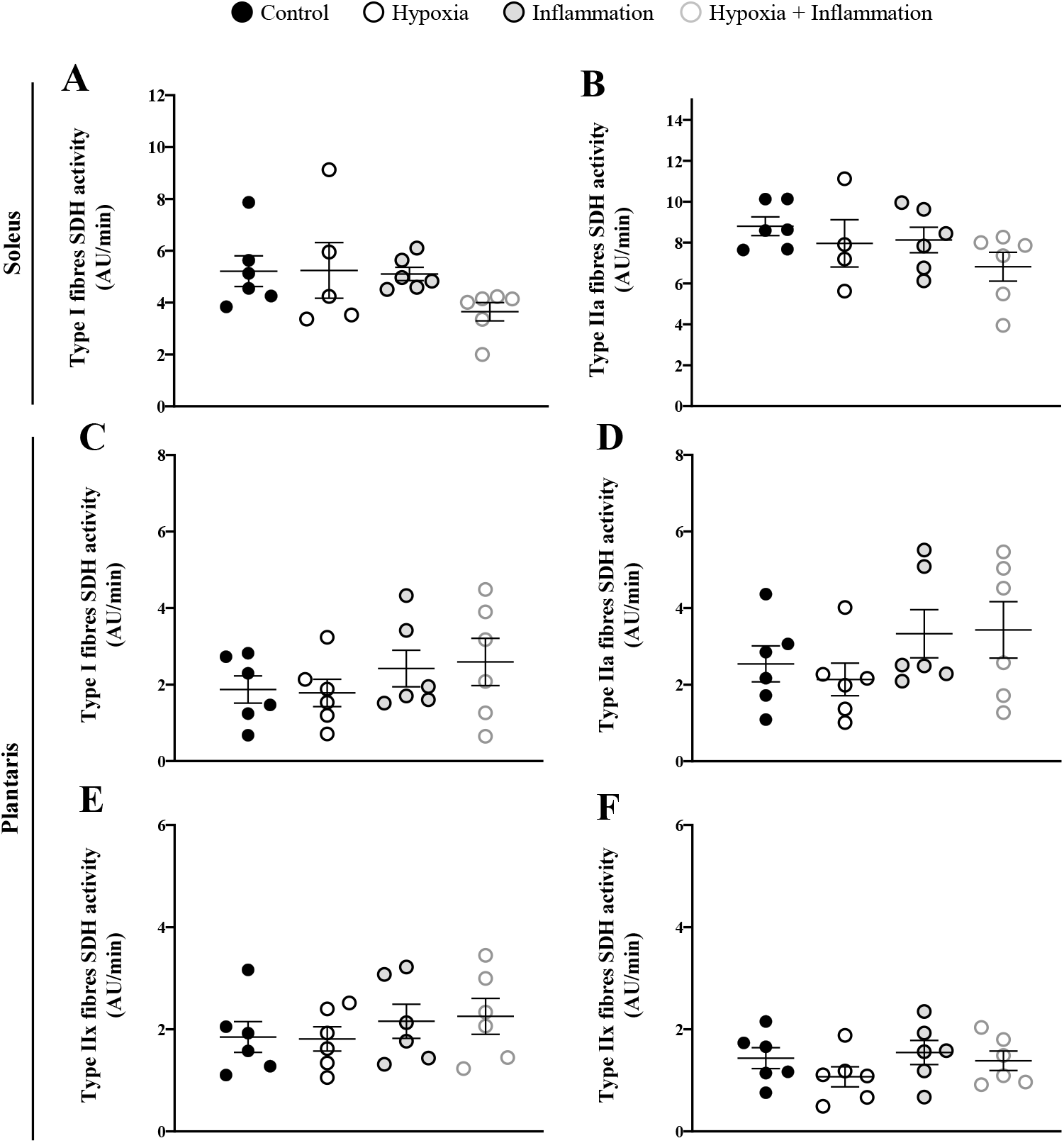
Effects of hypoxia and inflammation exposure on SDH activity of soleus and plantaris. SDH activity in type I (**A**) and type IIa (**B**) fibres of soleus and type I (**C**), type IIa (**D**), IIx (**E**) and type IIb (**F**) fibres of plantaris. Circles indicate individual data points and lines represent means ± SEM.

**Fig.8.**
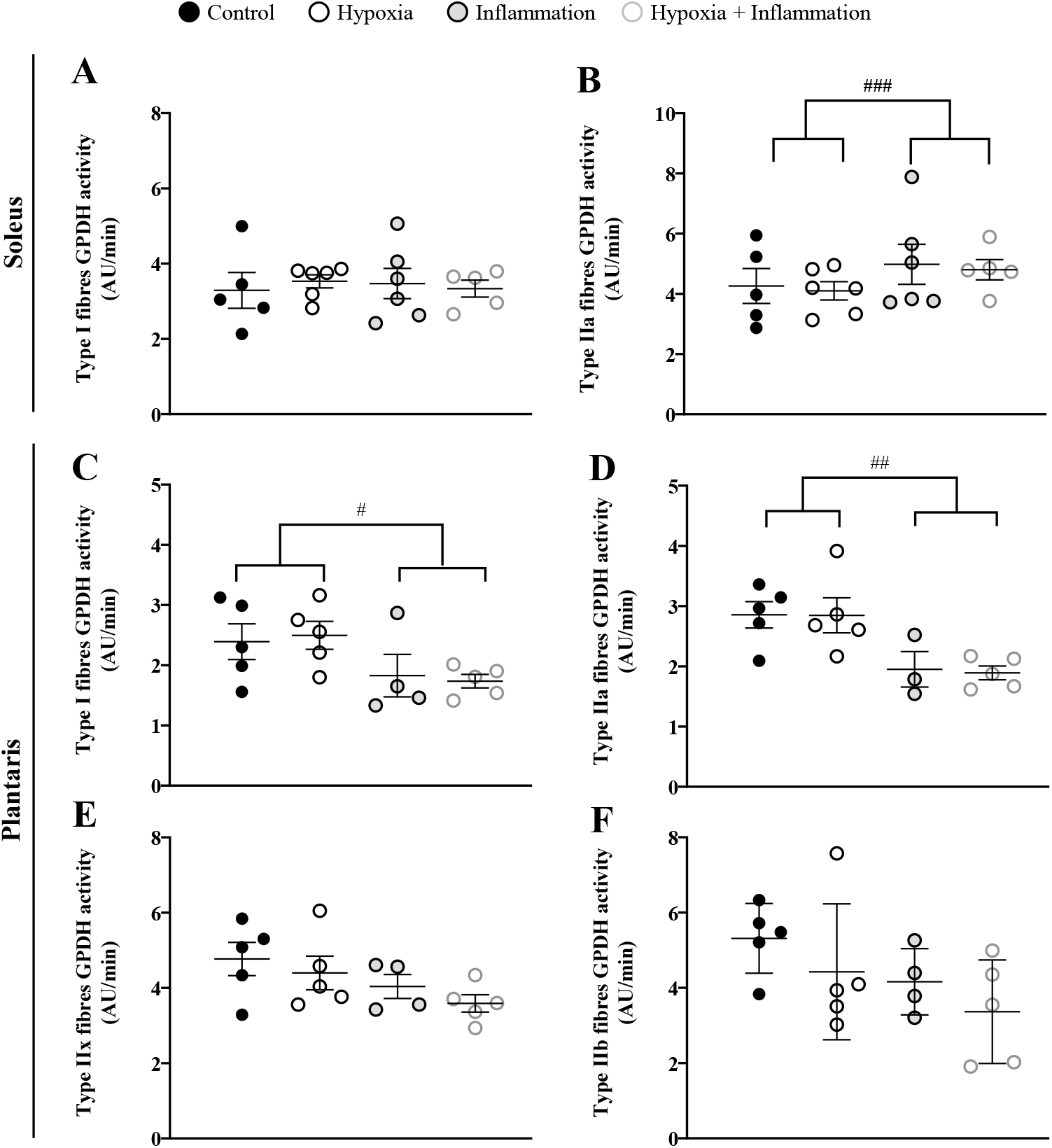
Effects of hypoxia and inflammation exposure on GPDH activity of soleus and plantaris. GPDH activity in type I (**A**) and type IIa (**B**) fibres of soleus and type I (**C**), type IIa (**D**), IIx (**E**) and type IIb (**F**) fibres of plantaris. Circles indicate individual data points and lines represent means ± SEM. **###:** p < 0.001, global effect of pulmonary inflammation (NC + H vs I + HI); **##:** p < 0.01, global effect of pulmonary inflammation (NC + H vs I + HI); **#:** p < 0.05, global effect of pulmonary inflammation (NC + H vs I + HI).

## 4. Discussion

Consistent with our hypothesis, chronic hypoxia impaired predominantly plantaris leading to mitochondrial dysfunction, evidenced by a decline in CI respiration and HAD activity, accompanied by a reduction in muscle weight and the CSA of type IIb fibres. Conversely, soleus exhibited greater resilience to hypoxia, preserving both its mass and respiratory properties, despite an elevated ROS emission. Chronic pulmonary inflammation triggered maladaptations in both muscles, as indicated by a dysregulation in mitochondrial calcium handling exclusively in soleus and divergent metabolic shifts in both soleus (a switch towards glycolysis in type IIa fibres) and plantaris (a lower reliance on glycolysis in type I and IIa fibres and a switch towards oxidative phosphorylation in type IIx and IIb fibres). Moreover, we have demonstrated that these alterations in skeletal muscle mitochondrial function and metabolic regulation can occur independently of major fibre type shifts.

### 4.1. Effects of chronic hypoxia

The reduction in ADP-stimulated CI respiration aligns with earlier studies demonstrating an impairment in mitochondrial respiratory capacity following chronic hypoxia (13,21). This may represent chronic adaptations to the reductive stress imposed by cellular hypoxia which leads to a buildup of reducing equivalents such as NADH and FADH_2_ and over-reduction of the inner mitochondrial membrane-embedded electron carriers (12). This redox imbalance can increase electron leak especially at the level of CI and CIII (12,22). As chronic hypoxia did not elicit any increase in plantaris overall or CI-derived H_2_O_2_ emission, it can be suggested that these changes represent a coordinated downregulation of oxidative metabolism to mitigate oxidative damage. A potential mechanism underlying this adaptation may partially involve the induction of NADH dehydrogenase ubiquinone 1 alpha subcomplex 4-like 2 (NDUFA4L2) by hypoxia-inducible factor 1α (HIF-1α) (23–25). Although we did not measure NDUFA4L2 expression, its induction under hypoxia has been demonstrated to dampen oxidative phosphorylation (OXPHOS) and ROS production in a wide variety of tissues including skeletal muscle (25), neonatal cardiomyocytes and brain tissue (23) as well as in hypoxic domains of cancer cells (24,26).

The hypoxia-induced decrease in plantaris HAD activity could reflect a diminished utilisation of fatty acids for ATP production and is consistent with past studies (27,28). This may be the result of an overall downregulation of all the mitochondrial pathways to prevent further provision of NADH and FADH_2_. We can also hypothesise about an inhibition of pyruvate dehydrogenase due to induction of pyruvate dehydrogenase kinase 1 by chronic hypoxia (29) that would thereby lower Krebs cycle activity and subsequently lead to less demand for Acetyl-CoA from fatty acids. This lower mitochondrial activity does not appear to be compensated by a higher glycolytic flux as hypoxia did not upregulate GPDH or LDH activity (supplementary Figure S7) in plantaris. Therefore, the energy availability might have been limiting with consequences on ATP-dependent processes such as protein synthesis. This is consistent with the reduction in plantaris weight and the CSA of type IIb fibres under hypoxia. Additionally, although we observed a decrease in food intake during the first 3 weeks of the hypoxic exposure, chronic hypoxia has been previously reported to exert hypophagia-independent muscle atrophy (14,18).

In contrast with plantaris, mitochondrial respiratory capacity and mass were preserved in soleus confirming that oxidative muscles are more resilient to chronic hypoxia. These findings are in the same line with earlier data demonstrating unchanged OXPHOS in soleus after 14 days at 10% FiO_2_ (30). Soleus postural role may necessitate sustained mitochondrial function which would also explain its mass preservation. However, the increase in CI-derived H_2_O_2_ emission appears to be a trade-off for preserving OXPHOS. To determine whether this rise stems from an intrinsic alteration in ROS generation or a proportional increase in CI basal respiration, H_2_O_2_ emission rates were normalised with the corresponding state 2 CI O_2_ consumption. Our results revealed a higher ratio in soleus under hypoxia, suggesting a greater fraction of electrons leaking to form ROS per O_2_. This observation may reflect a pathological ROS leak potentially explained by impaired H_2_O_2_ scavenging by catalase or glutathione peroxidase (31). Conversely, the unaltered mPTP sensitivity and the maintenance of mitochondrial respiration and muscle mass in hypoxic soleus suggest that this higher ROS production was without damaging effects. Although we did not assess manganese superoxide dismutase 2 (MnSOD2) activity, we can speculate that its activity may have increased to convert excess O_2_•-to the less reactive H_2_O_2_, representing a protective adaptation to attenuate mPTP and mitochondrial DNA damage.

### 4.2. Effects of chronic pulmonary inflammation

Although there was no effect of pulmonary inflammation on mitochondrial respiration of plantaris, we observed a downregulation in GPDH activity without any changes in SDH activity in its type I and type IIa fibres leading to a parallel higher SDH/GPDH. In contrast, given that there were no differences in the GPDH activity of type IIx and IIb fibres, their higher SDH/GPDH ratios suggests a shift towards oxidative metabolism. This is a novel finding implying that pulmonary inflammation may force the highly glycolytic fibres of plantaris to a metabolic state that is distant from its intrinsic functional capacity that may subsequently impact its peak power output. In addition, despite the increased reliance on oxidative metabolism, it is likely that the underdeveloped capillary network, lower levels of myoglobin and low levels of antioxidants defences observed in type IIx and IIb fibres (32) may structurally constrain their capacity to support OXPHOS.

In soleus, we observed a decrease in mitochondrial CRC following pulmonary inflammation indicating faster mPTP opening. This effect was prevented after inhibition of CyP-D by CsA, implying that this regulatory component of the mPTP may be directly modulated by pulmonary inflammation. Although we are the first to demonstrate increased mPTP sensitivity in skeletal muscle following pulmonary inflammation, comparable findings emanate from studies in muscle of COPD patients (6) or ageing muscles susceptible to chronic low-grade inflammation (33). The mPTP opening is known to be involved in pro-apoptotic factors release (34,35) followed by activation of proteolytic pathways and atrophy (36). Thus the absence of atrophy and decline in mitochondrial respiration may suggest that pulmonary inflammation provokes recurring, transient, low conductance pore openings that increases the sensitivity to Ca^2+^and constitutes a protective mechanism to reset soleus mitochondrial function. This alteration in calcium handling by soleus mitochondria may result from disrupted cytosolic Ca^2+^ homeostasis and sarcoplasmic reticulum (SR) stress initiated by proinflammatory mediators. For instance, Qaisar et al. (37) reported higher SR stress coupled with sarcoplasmic reticulum Ca^2+^-ATPase (SERCA) disruption in muscle of asthmatic patients. That promotes cytosolic Ca^2+^ accumulation and elevates mitochondrial Ca^2+^ uptake, leading to matrix Ca^2+^ overload, which sensitises the mPTP (38). Whether these recurrent low conductance openings may eventually transition to the more deleterious high-conductance mPTP openings (39) warrants further investigation.

Interestingly, we report that the SDH/GPDH ratio tended to decrease in type IIa fibres of soleus with pulmonary inflammation due to a higher GPDH activity suggesting a shift towards glycolysis that may signify detrimental functional consequences such as earlier fatigue during posture maintenance. Our findings align with increased hexokinase 2 mRNA expression in muscle of COPD patients presenting elevated tumour necrosis factor α (TNF-α) expression (40) and a glycolytic shift in myotubes exposed for 24 h to LPS (41). These alterations may stem from inflammation-induced impairments in mitochondrial biogenesis (40,42) and/or upregulation in mitophagic flux (43,44) resulting in reduced mitochondrial content in type IIa fibres and subsequent switch towards glycolysis.

### 4.3. Combined effect of chronic hypoxia and pulmonary inflammation

The original combination of hypoxia with pulmonary inflammation enabled us to investigate whether the simultaneous presence of these two stressors exacerbates skeletal muscle dysfunction beyond the effects of each stimulus alone. Our statistical analysis did not reveal any significant interactions suggesting that the combination of hypoxia and pulmonary inflammation does not result in additive effects that would compound muscle dysfunction. In contrast, our findings underscore that the two stimuli operate through distinct pathways that potentially compete with each other to elicit a response that is muscle- and parameter-specific. For instance, hypoxia appears to prevail over pulmonary inflammation with regards to plantaris state 3 CI respiration and HAD activity while pulmonary inflammation overrides hypoxia effects on soleus CRC. This likely reflects the opposing HIF-1α vs. TNF-α molecular signaling imposed by the two conditions and are consistent with previous work from our group showing distinct modulation of proteolysis and proteosynthesis pathways by hypoxia and pulmonary inflammation (17).

### 4.4. Strengths and Limitations

A major strength of our study was the simultaneous assessment of fibre typology and mitochondrial function in soleus and plantaris. Although skeletal muscle dysfunction in COPD has been proposed to stem from changes in fibre typology (20), we only report a rise in the percentage of plantaris IIx fibres in response to pulmonary inflammation. Since no other differences were observed, it is challenging to predict whether this rise in type IIx numbers represents a switch towards a more oxidative or glycolytic phenotype. Collectively, these results imply that mitochondrial intrinsic maladaptations may play a key role in the COPD-associated muscle dysfunction. A key strength of this study was the evaluation of the independent and combined effects of inflammation and hypoxia. We deliberately chose this model to address several objectives. Firstly, from a basic science standpoint, it enabled us to establish the unique fingerprint of each stressor, exploring whether the mitochondrial and metabolic responses are stress-dependent. Secondly, the study of their combined effects allowed us to decipher whether their interactions are synergistic, antagonistic or independent providing a new insight of the complex pathophysiology seen in these patients.

A limitation of our study is that we have not disentangled the proportion of metabolic and mitochondrial changes attributed to hypophagia associated with hypoxia. One would suggest the inclusion of a pair-fed NC group alongside the ad-libitum group. However, due to our simultaneous investigation of pulmonary inflammation that does not induce changes in food intake and the fact that hypoxia is intrinsically associated with alterations in food intake, we opted to feed the NC group ad-libitum and observe the global effects of hypoxia. Furthermore, while COPD being a disease mostly encountered in the ageing population (46), the use of young adult rats might have underestimated the magnitude of detrimental responses to hypoxia and pulmonary inflammation. Finally, only male rats were used and therefore our results are only applicable to male populations. Considering the sex differences in baseline mitochondrial bioenergetics (47) and antioxidant defences (48), future studies in female population should confirm our findings.

## 5. Conclusion

Using our original model of chronic hypoxia and pulmonary inflammation exposures, we showed muscle-specific mitochondrial and metabolic responses to those two key factors of COPD, with no overlap in their mechanism of action. Chronic hypoxia impairs predominantly the glycolytic-leaning plantaris compared to oxidative soleus while chronic pulmonary inflammation remodels the metabolic landscape of both muscles and increases the mPTP sensitivity in soleus. These alterations occur in the absence of fibre type shifts thereby suggesting that intrinsic mitochondrial defects may play a key role in the muscle dysfunction observed in COPD. Finally, distinct and opposing mechanisms of action between hypoxia and pulmonary inflammation highlight that the muscle response to these two stressors is not a simple sum of their isolated effects, but rather a complex interaction where one factor may modify or obscure the other and thus targeted interventions for hypoxic versus inflammatory muscle defects should be developed.

## Supporting information

Supplemental datas

## 6. Supplementary information

### Supplemental data

The supplementary figures are available in a separate file.

## 7. Acknowledgments

A.G has been supported by a 3-year Ph.D. fellowship by the foundation: ‘Agir pour les maladies chroniques’. We acknowledge the LBFA Imaging facility, member of Grenoble-IsDV IBISA labeled perimeter (Grenoble Imagerie Sciences du Vivant) and member of the national infrastructure France-BioImaging supported by the French National Research Agency (ANR-10-INBS-04-01).

## References

1. Taivassalo T, Hepple RT. Integrating Mechanisms of Exacerbated Atrophy and Other Adverse Skeletal Muscle Impact in COPD. Front Physiol. 2022 June 3;13:861617.

2. Gosker HR, Hesselink MKC, Duimel H, Ward KA, Schols AMWJ. Reduced mitochondrial density in the vastus lateralis muscle of patients with COPD. Eur Respir J. 2007 July;30(x1):73–9.

3. Puente-Maestu L, Pérez-Parra J, Godoy R, Moreno N, Tejedor A, González-Aragoneses F, et al. Abnormal mitochondrial function in locomotor and respiratory muscles of COPD patients. Eur Respir J. 2009 May;33(5):1045–52.

4. Maltais F. Oxidative enzyme activities of the vastus lateralis muscle and the functional status in patients with COPD. Thorax. 2000 Oct 1;55(10):848–53.

5. Van Den Borst B, Slot IGM, Hellwig VACV, Vosse BAH, Kelders MCJM, Barreiro E, et al. Loss of quadriceps muscle oxidative phenotype and decreased endurance in patients with mild-to-moderate COPD. J Appl Physiol. 2013 May 1;114(9):1319–28.

6. Puente-Maestu L, Pérez-Parra J, Godoy R, Moreno N, Tejedor A, Torres A, et al. Abnormal Transition Pore Kinetics and Cytochrome C Release in Muscle Mitochondria of Patients with Chronic Obstructive Pulmonary Disease. Am J Respir Cell Mol Biol. 2009 June;40(6):746–50.

7. Gea J, Pascual S, Casadevall C, Orozco-Levi M, Barreiro E. Muscle dysfunction in chronic obstructive pulmonary disease: update on causes and biological findings. J Thorac Dis. 2015;7(10).

8. Jaitovich A, Barreiro E. Skeletal Muscle Dysfunction in Chronic Obstructive Pulmonary Disease. What We Know and Can Do for Our Patients. Am J Respir Crit Care Med. 2018 July 15;198(2):175–86.

9. Barnes PJ, Celli BR. Systemic manifestations and comorbidities of COPD. Eur Respir J. 2009 May;33(5):1165–85.

10. Eliason G, Abdel-Halim SM, Piehl-Aulin K, Kadi F. Alterations in the muscle-to-capillary interface in patients with different degrees of chronic obstructive pulmonary disease. Respir Res. 2010 Dec;11(1):97.

11. McNicholas W, Kent, Mitchell. Hypoxemia in patients with COPD: cause, effects, and disease progression. Int J Chron Obstruct Pulmon Dis. 2011 Mar;199.

12. Clanton TL. Hypoxia-induced reactive oxygen species formation in skeletal muscle. J Appl Physiol. 2007 June;102(6):2379–88.

13. Gamboa JL, Andrade FH. Muscle endurance and mitochondrial function after chronic normobaric hypoxia: contrast of respiratory and limb muscles. Pflüg Arch - Eur J Physiol. 2012 Feb;463(2):327–38.

14. Favier FB, Britto FA, Freyssenet DG, Bigard XA, Benoit H. HIF-1-driven skeletal muscle adaptations to chronic hypoxia: molecular insights into muscle physiology. Cell Mol Life Sci. 2015 Dec;72(24):4681–96.

15. Hoppeler H, Vogt M, Weibel ER, Flück M. Response of Skeletal Muscle Mitochondria to Hypoxia. Exp Physiol. 2003 Jan;88(1):109–19.

16. Ji Y, Li M, Chang M, Liu R, Qiu J, Wang K, et al. Inflammation: Roles in Skeletal Muscle Atrophy. Antioxidants. 2022 Aug 29;11(9):1686.

17. Chabert C, Khochbin S, Rousseaux S, Furze R, Smithers N, Prinjha R, et al. Muscle hypertrophy in hypoxia with inflammation is controlled by bromodomain and extra-terminal domain proteins. Sci Rep. 2017 Sept 21;7(1):12133.

18. De Theije CC, Langen RCJ, Lamers WH, Gosker HR, Schols AMWJ, Köhler SE. Differential sensitivity of oxidative and glycolytic muscles to hypoxia-induced muscle atrophy. J Appl Physiol. 2015 Jan 15;118(2):200–11.

19. Picard M, Hepple RT, Burelle Y. Mitochondrial functional specialization in glycolytic and oxidative muscle fibers: tailoring the organelle for optimal function. Am J Physiol-Cell Physiol. 2012 Feb 15;302(4):C629– 41.

20. Hartmann JP, Caldwell HG, Iepsen UW. Muscle fibre type shift in COPD: Adaptive, maladaptive or a bit of both? Exp Physiol. 2025 Mar 3;EP092126.

21. Jacobs RA, Boushel R, Wright-Paradis C, Calbet JAL, Robach P, Gnaiger E, et al. Mitochondrial function in human skeletal muscle following high-altitude exposure. Exp Physiol. 2013 Jan;98(1):245–55.

22. Murphy MP. How mitochondria produce reactive oxygen species. Biochem J. 2009 Jan 1;417(1):1–13.

23. Tello D, Balsa E, Acosta-Iborra B, Fuertes-Yebra E, Elorza A, Ordóñez Á, et al. Induction of the Mitochondrial NDUFA4L2 Protein by HIF-1α Decreases Oxygen Consumption by Inhibiting Complex I Activity. Cell Metab. 2011 Dec;14(6):768–79.

24. Lai RKH, Xu IMJ, Chiu DKC, Tse APW, Wei LL, Law CT, et al. NDUFA4L2 Fine-tunes Oxidative Stress in Hepatocellular Carcinoma. Clin Cancer Res. 2016 June 15;22(12):3105–17.

25. Liu Z, Chaillou T, Santos Alves E, Mader T, Jude B, Ferreira DMS, et al. Mitochondrial NDUFA4L2 is a novel regulator of skeletal muscle mass and force. FASEB J [Internet]. 2021 Dec [cited 2025 Sept 25];35(12). Available from: https://onlinelibrary.wiley.com/doi/10.1096/fj.202100066R

26. Minton DR, Fu L, Mongan NP, Shevchuk MM, Nanus DM, Gudas LJ. Role of NADH Dehydrogenase (Ubiquinone) 1 Alpha Subcomplex 4-Like 2 in Clear Cell Renal Cell Carcinoma. Clin Cancer Res. 2016 June 1;22(11):2791–801.

27. Galbès O, Goret L, Caillaud C, Mercier J, Obert P, Candau R, et al. Combined effects of hypoxia and endurance training on lipid metabolism in rat skeletal muscle. Acta Physiol. 2008 June;193(2):163–73.

28. Levett DZ, Radford EJ, Menassa DA, Graber EF, Morash AJ, Hoppeler H, et al. Acclimatization of skeletal muscle mitochondria to high-altitude hypoxia during an ascent of Everest. FASEB J. 2012 Apr;26(4):1431– 41.

29. De Palma S, Ripamonti M, Viganò A, Moriggi M, Capitanio D, Samaja M, et al. Metabolic Modulation Induced by Chronic Hypoxia in Rats Using a Comparative Proteomic Analysis of Skeletal Muscle Tissue. J Proteome Res. 2007 May 1;6(5):1974–84.

30. Horscroft JA, Burgess SL, Hu Y, Murray AJ. Altered Oxygen Utilisation in Rat Left Ventricle and Soleus after 14 Days, but Not 2 Days, of Environmental Hypoxia. Sturmey R, editor. PLOS ONE. 2015 Sept 21;10(9):e0138564.

31. Sullivan-Gunn MJ, Lewandowski PA. Elevated hydrogen peroxide and decreased catalase and glutathione peroxidase protection are associated with aging sarcopenia. BMC Geriatr. 2013 Dec;13(1):104.

32. Vasileiadou O, Nastos GG, Chatzinikolaou PN, Papoutsis D, Vrampa DI, Methenitis S, et al. Redox Profile of Skeletal Muscles: Implications for Research Design and Interpretation. Antioxidants. 2023 Sept 7;12(9):1738.

33. Gouspillou G, Sgarioto N, Kapchinsky S, Purves-Smith F, Norris B, Pion CH, et al. Increased sensitivity to mitochondrial permeability transition and myonuclear translocation of endonuclease G in atrophied muscle of physically active older humans. FASEB J. 2014 Apr;28(4):1621–33.

34. Endlicher R, Drahota Z, Štefková K, Červinková Z, KuČera O. The Mitochondrial Permeability Transition Pore—Current Knowledge of Its Structure, Function, and Regulation, and Optimized Methods for Evaluating Its Functional State. Cells. 2023 Apr 27;12(9):1273.

35. Carraro M, Bernardi P. The mitochondrial permeability transition pore in Ca2+ homeostasis. Cell Calcium. 2023 May;111:102719.

36. Li A, Yi J, Li X, Zhou J. Physiological Ca2+ Transients Versus Pathological Steady-State Ca2+ Elevation, Who Flips the ROS Coin in Skeletal Muscle Mitochondria. Front Physiol. 2020 Oct 22;11:595800.

37. Qaisar R, Qayum M, Muhammad T. Reduced sarcoplasmic reticulum Ca2+ ATPase activity underlies skeletal muscle wasting in asthma. Life Sci. 2021 May;273:119296.

38. Zhou J, Dhakal K, Yi J. Mitochondrial Ca2+ uptake in skeletal muscle health and disease. Sci China Life Sci. 2016 Aug;59(8):770–6.

39. Korge P, Yang L, Yang JH, Wang Y, Qu Z, Weiss JN. Protective Role of Transient Pore Openings in Calcium Handling by Cardiac Mitochondria. J Biol Chem. 2011 Oct;286(40):34851–7.

40. Remels AH, Schrauwen P, Broekhuizen R, Willems J, Kersten S, Gosker HR, et al. Peroxisome proliferator-activated receptor expression is reduced in skeletal muscle in COPD. Eur Respir J. 2007 Aug;30(2):245–52.

