## Supplemental datas for "Skeletal muscle fibre type determines mitochondrial and metabolic responses to hypoxia and pulmonary inflammation"

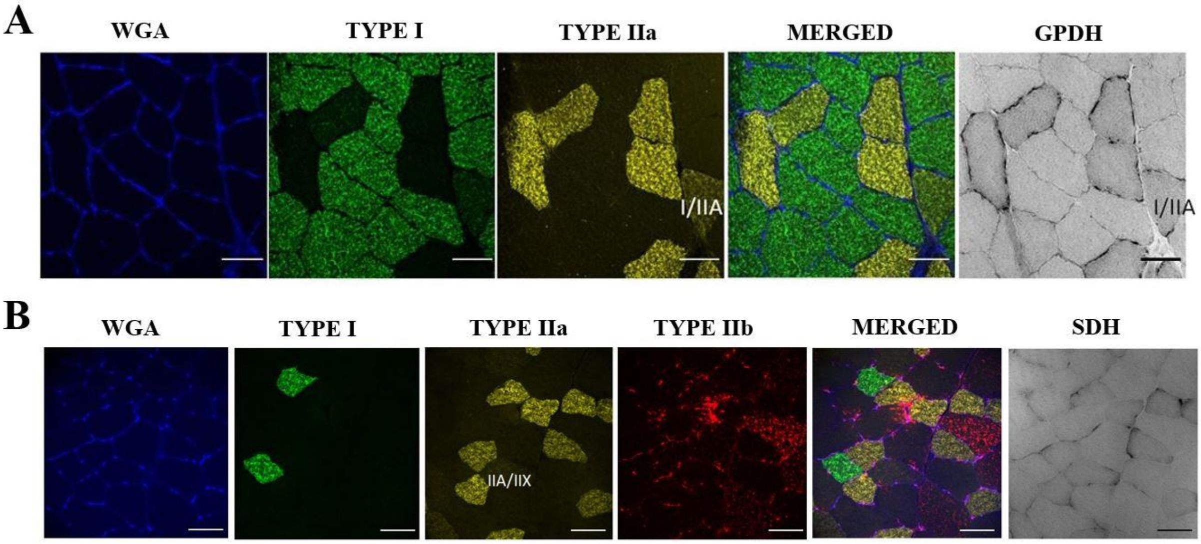


***Fig.S1*. Representative images of MHC, WGA, GPDH and SDH staining.** *Images show WGA, MHC I, MHC IIA, merged (WGA + MHC I + MHC IIa) and GPDH staining for soleus* ***(A)*** *and WGA, MHC I, MHC IIa, MHC IIb, merged (WGA + MHC I + MHC IIa + MHC IIb) and SDH staining for plantaris* ***(B)****. Scale bar, 50 µm.*

SUPPLEMENTARY FIGURES

Skeletal muscle fibre type determines mitochondrial and metabolic responses to hypoxia and pulmonary inflammation

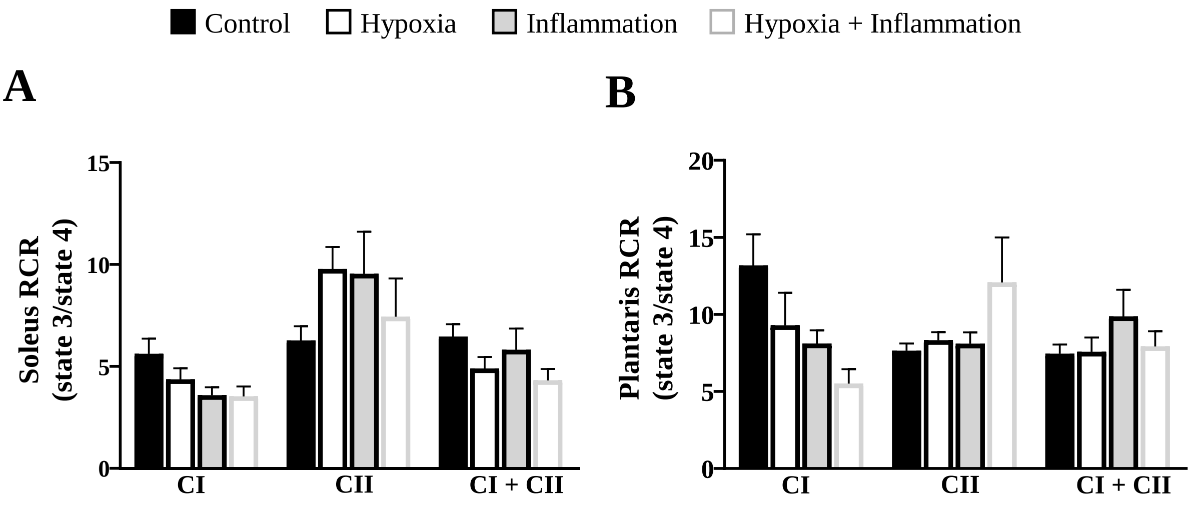


***Fig.S2. Respiratory control ratio in isolated mitochondria from soleus and plantaris muscle after exposure to hypoxia and inflammation****. Soleus* ***(A)*** *and plantaris* ***(B)*** *RCR (state 3/state 4) with CI (GM), CII (S) and CI + CII (GMS) substrates. Data are shown as mean (bars) ± SEM (error bars).*

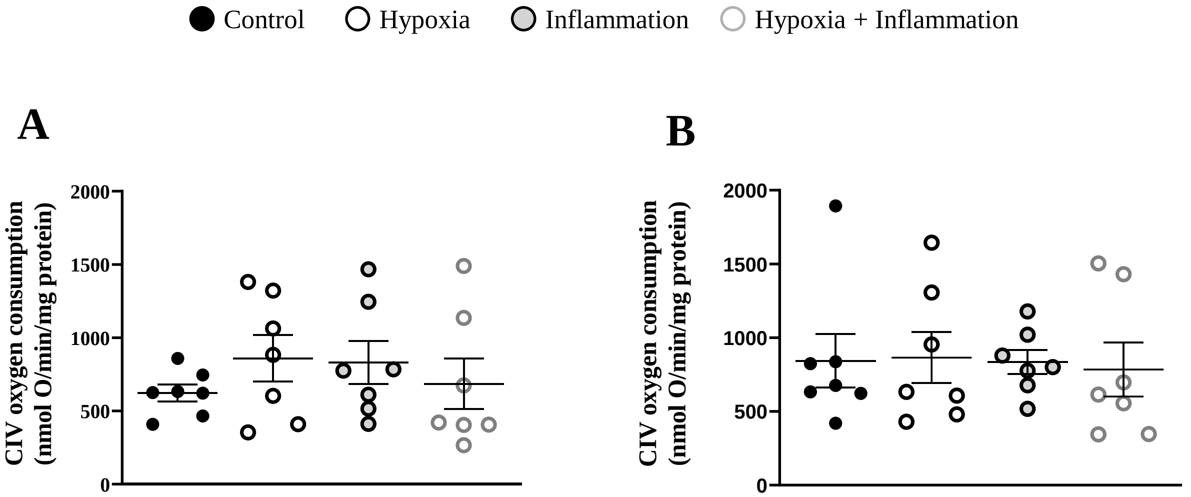


***Fig.S3. Complex IV oxygen consumption in isolated mitochondria from soleus and plantaris muscle after exposure to hypoxia and inflammation.*** *Soleus* ***(A)*** *and plantaris* ***(B)*** *uncoupled mitochondrial respiration measured by oxygraphy using AA, ascorbate, TMPD and DNP. Circles indicate individual data points and lines represent means ± SEM.*

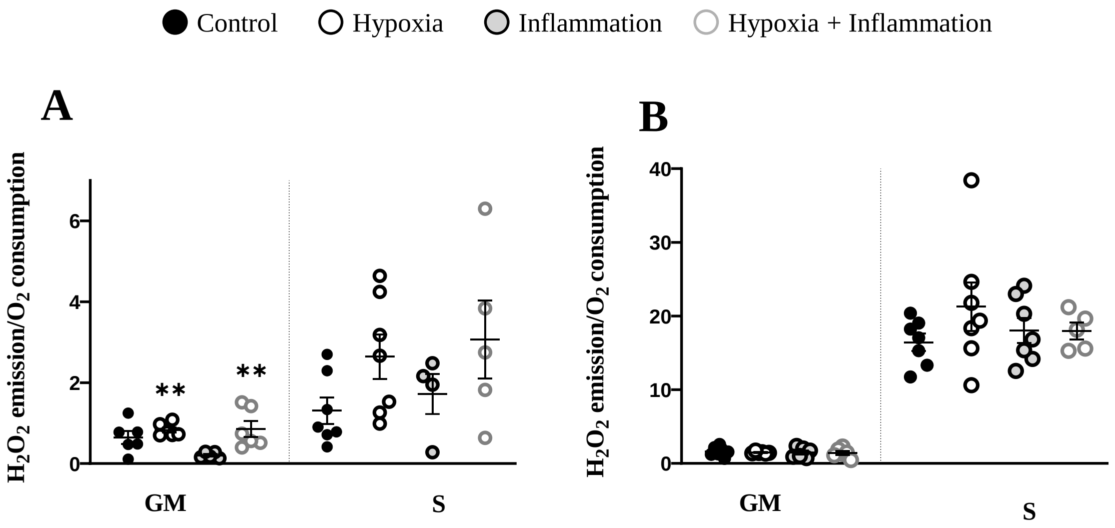


***Fig.S4. Soleus (A) and plantaris (B) mitochondria $H_2O_2$ release with GM or S were normalised to CI or CII basal (substrate only) oxygen consumption, respectively.*** *Circles indicate individual and lines represent means ± SEM. ** $p < 0.01$, global effect of hypoxia (NC + I vs H + HI).*

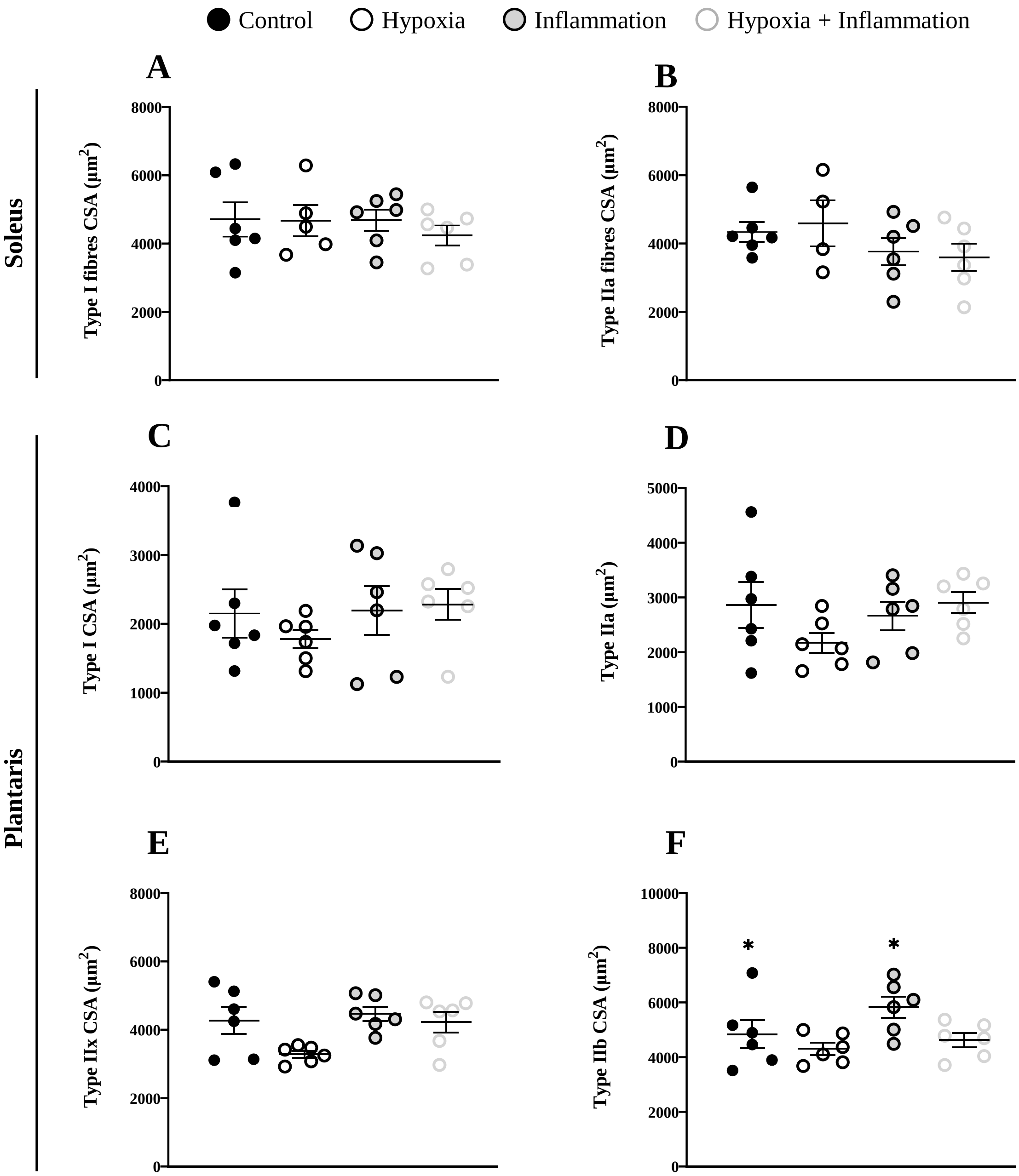


***Fig.S5. Effects of hypoxia and inflammation exposure on fibre cross sectional area.*** *CSA analysis in type I **(A)** and type IIa **(B)** fibres of soleus and type I **(C)**, type IIa **(D)**, IIx **(E)** and type IIb **(F)** fibres of plantaris. Circles indicate individual data points and lines represent means ± SEM. *: p < 0.05, global effect of hypoxia (NC + I vs H + HI)*

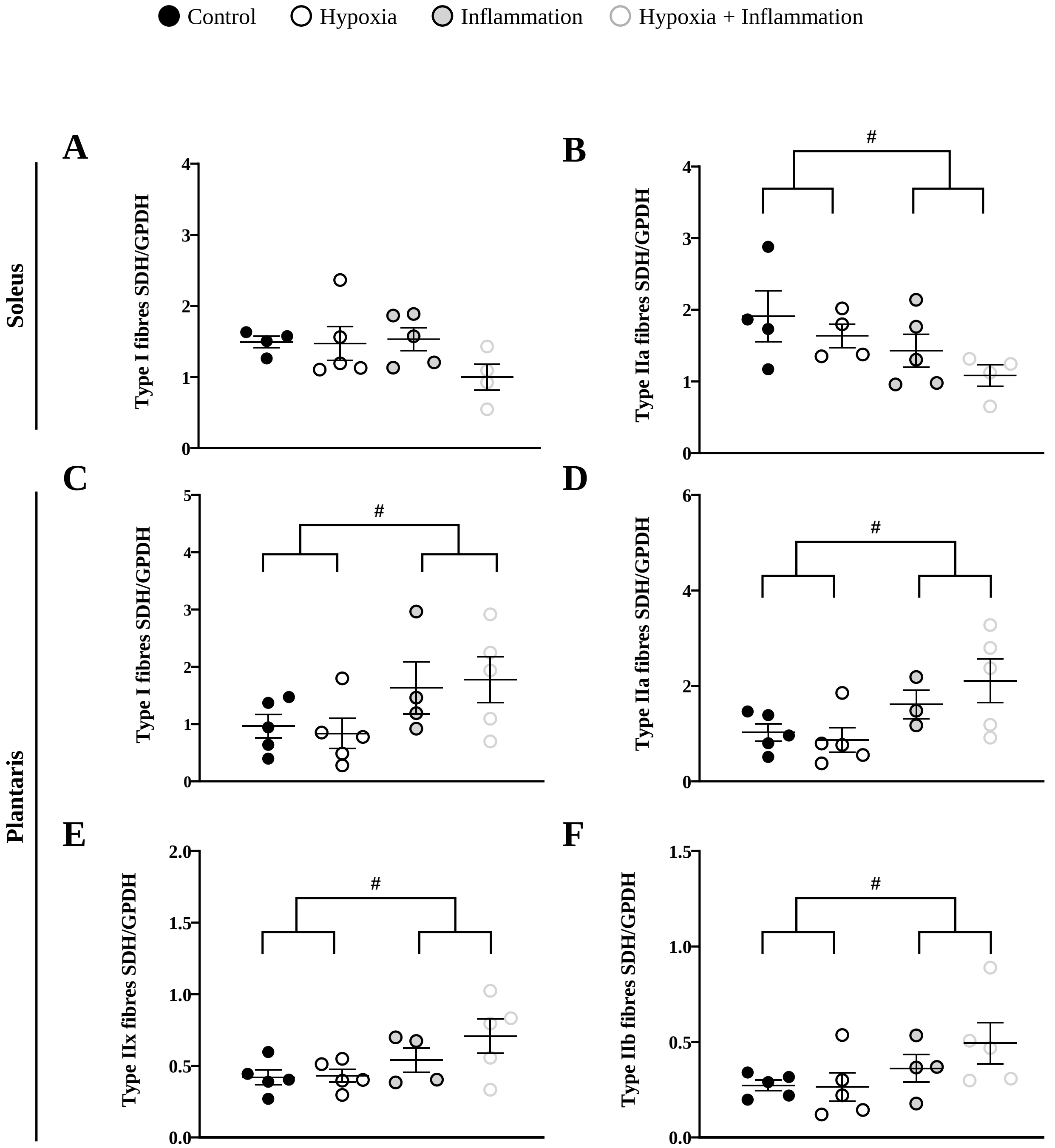


***Fig.S6 Effects of hypoxia and inflammation exposure on SDH/GPDH activity ratio of soleus and plantaris.*** *SDH/GPDH ratio in type I (**A**) and type IIa (**B**) fibres of soleus and type I (**C**), type IIa (**D**), IIx (**E**) and type IIb (**F**) fibres of plantaris. Circles indicate individual data points and lines represent means ± SEM.* ***#:*** *$p < 0.05$, global effect of pulmonary inflammation (NC + H vs I + HI).*

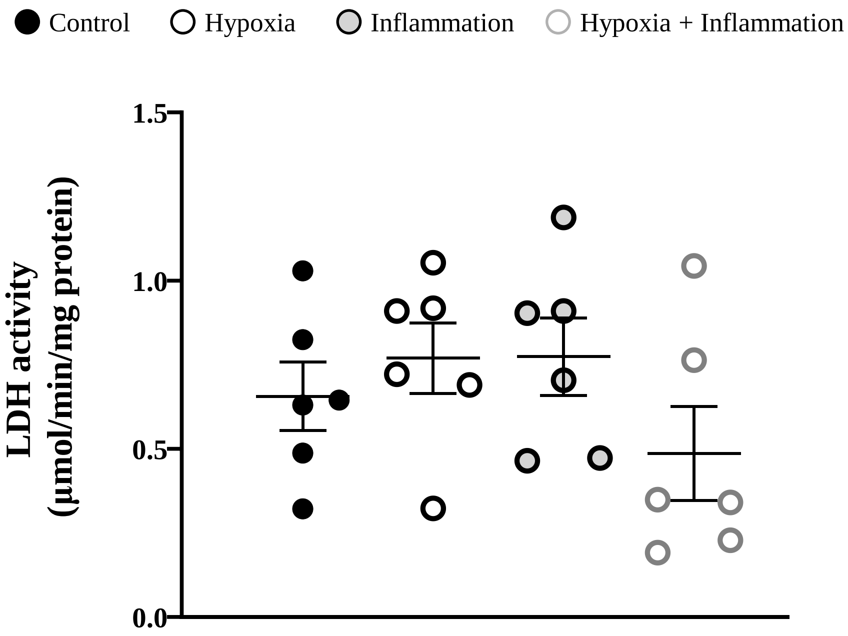


***Fig.S7. Effect of hypoxia and inflammation exposure on plantaris LDH activity.*** *LDH activity was measured in the cytosolic fraction of plantaris. Circles indicate individual data points and lines represent means ± SEM.*

### SUPPLEMENTARY DATA REFERENCE LIST

41. Eggelbusch M, Shi A, Broeksma BC, Vázquez-Cruz M, Soares MN, De Wit GMJ, et al. The NLRP3 inflammasome contributes to inflammation-induced morphological and metabolic alterations in skeletal muscle. J Cachexia Sarcopenia Muscle. 2022 Dec;13(6):3048–61.

42. Remels AHV, Gosker HR, Schrauwen P, Hommelberg PPH, Sliwinski P, Polkey M, et al. TNF-α impairs regulation of muscle oxidative phenotype: implications for cachexia? FASEB J. 2010 Dec;24(12):5052–62.

43. Leermakers PA, Remels AHV, Langen RCJ, Schols AMWJ, Gosker HR. Pulmonary inflammation-induced alterations in key regulators of mitophagy and mitochondrial biogenesis in murine skeletal muscle. BMC Pulm Med. 2020 Dec;20(1):20.

44. Leermakers PA, Schols AMWJ, Kneppers AEM, Kelders MCJM, De Theije CC, Lainscak M, et al. Molecular signalling towards mitochondrial breakdown is enhanced in skeletal muscle of patients with chronic obstructive pulmonary disease (COPD). Sci Rep. 2018 Oct 9;8(1):15007.

45. Li X, Berg NK, Mills T, Zhang K, Eltzschig HK, Yuan X. Adenosine at the Interphase of Hypoxia and Inflammation in Lung Injury. Front Immunol. 2021 Jan 14;11:604944.

46. Jarhyan P, Hutchinson A, Khaw D, Prabhakaran D, Mohan S. Prevalence of chronic obstructive pulmonary disease and chronic bronchitis in eight countries: a systematic review and meta-analysis. Bull World Health Organ. 2022 Mar 1;100(03):216–30.

47. Miotto PM, McGlory C, Holloway TM, Phillips SM, Holloway GP. Sex differences in mitochondrial respiratory function in human skeletal muscle. Am J Physiol-Regul Integr Comp Physiol. 2018 June 1;314(6):R909–15.

48. Tiberi J, Cesarini V, Stefanelli R, Canterini S, Fiorenza MT, La Rosa P. Sex differences in antioxidant defence and the regulation of redox homeostasis in physiology and pathology. Mech Ageing Dev. 2023 Apr;211:111802.
